# The Transcription Factor OsNAC5 Mediates Cadmium Stress Response and Accumulation in Rice

**DOI:** 10.1101/2024.04.07.588483

**Authors:** Shubao Hu, Jinfen Chen, Hui Wang, E Ji, Xinxin Su, Muyao Zhu, Xiaoyan Xiang, Li Gong, Qiang Zhou, Xin Xiao, Ganlin Wu, Hannie Zha

## Abstract

Cadmium (Cd) is a toxic heavy metal that endangers human health through contaminated rice consumption. Reducing Cd uptake in rice is essential for minimizing grain Cd accumulation. However, the transcriptional regulation of Cd uptake in rice remains largely elusive. This study reveals that the transcription factor OsNAC5 in *Oryza sativa* positively regulates the Cd transporter gene *OsNRAMP1*, influencing Cd uptake. OsNAC5 is primarily expressed in roots, localized to the nucleus, and upregulated by Cd-induced H_2_O_2_. Knocking out *OsNAC5* reduces Cd levels in both shoots and roots and increases Cd sensitivity. *OsNRAMP1* expression, enhanced by Cd stress, depends on OsNAC5, which binds to “CATGTG” motifs in the *OsNRAMP1* promoter activating its expression. Loss of *OsNAC5* function decreases Cd accumulation in rice grains. Our findings elucidate the transcriptional regulation of Cd stress response in rice and suggest biotechnological approaches to reduce Cd uptake in crops.

**Declaration of Competing Interest:** The authors declare that they have no known competing financial interests or personal relationships that could have appeared to influence the work reported in this paper.

## 1. Introduction

Cadmium (Cd) is a highly toxic heavy metal that poses a threat to human health, as it can cause kidney damage and increase the risk of cancer (Nordberg, 2009). Environmental contamination with Cd arises from various anthropogenic activities, including mining, the use of Cd-containing fertilizers, and the application of sewage sludge (Zhao et al., 2015). Rice, a staple crop in many Asian countries, exhibits higher Cd uptake and accumulation compared to other cereals such as wheat and maize (Sui et al., 2018). The current levels of Cd exposure in most populations exceed the threshold for adverse health effects, underscoring the urgent need for strategies to mitigate Cd transfer from soil to edible plant parts and protect human health (Chen et al., 2018).

Recent advances have been made in identifying key transporters involved in Cd uptake and translocation in plants (Clemens and Ma, 2016). The natural resistance-associated macrophage protein (NRAMP) family, particularly OsNRAMP5, has been identified as a major influx transporter for manganese (Mn), Cd and Lead (Pb) in rice (Chang et al., 2022; Ishikawa et al., 2012; Ishimaru et al., 2012; Sasaki et al., 2012; Yang et al., 2014). OsNRAMP5 is localized at the plasma membrane of the exodermis and endodermis in the rice roots, facilitating the uptake of these metals (Chang et al., 2022; Sasaki et al., 2012). However, mutations in OsNRAMP5, while effective in restricting Cd uptake, result in compromised growth and grain yield under Mn-limiting conditions (Sasaki et al., 2012; Yang et al., 2014). Conversely, overexpression of *OsNRAMP5* has been shown to decrease Cd accumulation in the shoots and grains by disrupting radial Cd transport into the stele (Chang et al., 2020). OsNRAMP1, another member of the NRAMP family, shares high amino acid sequence similarity with OsNRAMP5 and has been implicated in Mn and Cd uptake in rice (Chang et al., 2020; Takahashi et al., 2011). Both OsNRAMP1 and OsNRAMP5 contribute to metal uptake, although their functions are not completely redundant (Chang et al., 2020; Takahashi et al., 2011). Once internalized, Cd is transported to vacuoles by OsHMA3, a P-type heavy metal ATPase, or translocated to shoots via OsHMA2 (Ueno et al., 2010). Despite the understanding of these transporters’ roles, the transcriptional regulatory mechanisms controlling their expression are not well characterized.

However, the transcriptional regulatory mechanisms controlling Cd accumulation in crops remain poorly understood due to a lack of research in this area. Overexpression of *OsNAC5* in transgenic plants has been shown to provide increased tolerance to high salinity (Takasaki et al., 2010). As a transcriptional activator, OsNAC5 enhances stress tolerance by increasing the expression of stress-responsive genes in rice, such as late embryogenesis abundant 3 (*OsLEA3*) (Takasaki et al., 2010). Song et al. (2011) reported that OsNAC5 plays a significant role in enhancing tolerance to abiotic stress in rice by modulating downstream targets associated with the accumulation of compatible solutes, sodium ions (Na^+^), hydrogen peroxide (H_2_O_2_), and malondialdehyde (MDA). Jeong et al. (2013) showed that root-specific overexpression of *OsNAC5* leads to substantial root enlargement, which in turn improves drought tolerance and increases grain yield under field conditions. However, the roles of OsNAC5 in Cd tolerance and accumulation have yet to be fully explored.

In this study, we aimed to investigate the role of OsNAC5 in the regulation of Cd tolerance and uptake in rice. Through analysis of its expression pattern, subcellular localization, phenotypic response, and Cd concentration, we provide compelling evidence that OsNAC5 significantly influences Cd tolerance and accumulation in rice. Our findings shed light on the importance of OsNAC5 in the response of rice to Cd stress, highlighting its potential as a target for improving Cd tolerance in rice crops. Moreover, we demonstrate that OsNAC5 directly interacts with “ CATGTG” motifs in the promoter of OsNRAMP1, promoting its expression.

## 2. Materials and Methods

### 2.1. Generation of rice genetic materials and plant growth conditions

Generation of The *osnac5* Knockout Line: The *osnac5* knockout line was generated using the CRISPR/Cas9 genome editing system in the ‘ZH11’ rice cultivar background. A specific guide RNA (gRNA) sequence (297–319 bp) was used for this purpose (Supplementary Table S1). Agrobacterium tumefaciens-mediated transformation was performed on actively dividing calli derived from mature rice embryos (Nishimura et al., 2006). Homozygous mutant lines were confirmed by sequencing PCR-amplified genomic fragments from putative mutants.

Construction of Genetic Complementation Lines: For the genetic complementation of the *osnac5* mutant, a genomic fragment encompassing the *OsNAC5* promoter (1011 bp upstream of the start codon) and codon sequence (990 bp starting from ‘ATG’) was PCR-amplified from rice root DNA. This fragment was subsequently cloned into the binary vector pCAMBIA1300-HA.The primers used for amplification are listed in Supplementary Table S1. The resulting construct, ProOsNAC5::OsNAC5-HA, was introduced into the *osnac5* mutant plants (free of selection marker) via Agrobacterium-mediated transformation. Two independent complementation lines (ProOsNAC5::OsNAC5-HA, T3 generation) were selected based on immunoblotting analysis using anti-HA antibody. These lines are used for further phenotypic characterization and chromatin immunoprecipitation (ChIP) assays.

Growth Conditions and Nutrient Solution: Seeds of both wild-type ‘ZH11’ and *osnac5* mutant rice were initially soaked in water for 48 hours, followed by germination in a 0.5 mM CaCl_2_ solution. After five days, seedlings were transferred to a half-strength Kimura B solution (pH 5.5) and grown in a greenhouse with controlled conditions at 25—30 ℃. The nutrient solution was composed of 0.18 mM (NH_4_)_2_SO_4_, 0.27 mM MgSO_4_·7H_2_O, 0.09 mM KNO_3_, 0.18 mM Ca(NO_3_)_2_·4H_2_O, 0.09 mM KH_2_PO_4_, 0.5 µM MnCl_2_·4H_2_O, 3 µM H_3_BO_3_, 1 µM (NH_4_)_6_Mo_7_O_24_·4H_2_O, 0.4 µM ZnSO_4_·7H_2_O, 0.2 µM; CuSO_4_·5H_2_O, and 100 µM Fe(III)-EDTA. This solution was refreshed every two days to ensure optimal nutrient availability.

### 2.2. Subcellular localization

A cDNA fragment encoding *OsNAC5* was amplified from a rice cDNA library using OsNAC5-specific primers (Supplementary Table S1.). The PCR product was cloned into the pBluescript SK(+) vector, which harbors the 35S cauliflower mosaic virus (CaMV) promoter, in order to drive constitutive expression. For subcellular localization studies, *OsNAC5* coding sequence was fused in-frame with the coding sequence of Yellow Fluorescent Protein (YFP) fusion protein to generate an OsNAC5-YFP fusion construct. This expression cassette was then introduced into rice protoplasts using the polyethylene glycol (PEG) 3350/Ca^2+^ transfection method (Shi et al., 2014). Following transfection, the rice protoplasts were incubated in the dark at 25 ℃ for 12–18 hours to allow for expression of the fusion protein. After the incubation period, the rice protoplasts expressing the OsNAC5-YFP fusion protein were examined for fluorescence localization. Images were captured using a laser confocal microscope (UltraVIEW VOX, PerkinElmer, Waltham, MA, USA).

### 2.3. Gene expression analysis

To elucidate the tissue-specific expression pattern of *OsNAC5* in wild-type (WT) rice, samples were harvested from various tissues, including roots, basal stems, mature leaf blades, mature leaf sheaths, young leaf blades, young leaf sheaths, of two-week-old seedlings. To evaluate the expression of *OsNAC5* under Cadmium Nitrate (Cd(NO_3_)_2_) stress, WT rice seedlings at the same developmental stage were subjected to a gradient of Cd(NO_3_)_2_ concentration (0 to 100 µM) and to hydrogen peroxide (H_2_O_2_, 10 mM) for predetermined time periods. Furthermore, to investigate the expression profiles of Cd transporter genes (*OsNRAMP1*, *OsNRAMP5*, *OsHMA2*, and *OsHMA3*), as well as other potential downstream functional genes (*OsLEA3*, *OsNAS3*, and *OsYSL2*) of *OsNAC5*, 2-week-old WT rice seedlings were exposed to 10 µM Cd(NO_3_)_2_ for a duration of 12 hours.

For RNA extraction, total RNA was extracted from the treated rice tissues using a Fastpure Plant Total RNA Isolation Mini Kit (Product code: RC401, Vazyme, Nanjing, China). First-strand cDNA synthesis was performed using the HiScript II 1st Strand cDNA Synthesis Kit (Product code: R211-01, Vazyme) with 1 µg of the extracted total RNA as a template. RT-qPCR analysis was performed using the ChamQ SYBR Color qPCR Master Mix (Product code: Q311-02, Vazyme), with *Actin* being utilized as the internal control for normalization. Primer sequences for gene expression analysis are detailed in Supplementary Table S1.

### 2.4. Phenotypic analysis

To evaluate the impact of Cd stress on plant phenotype, seeds from wild-type (WT) rice, the *osnac5* mutant, and two independent *osnac5* complementation lines were germinated. Post-germination, seedlings were placed on nets floating in 4-liter containers of solution. Two-week-old seedlings were then exposed to 0, 10, and 100 µM Cd(NO_3_)_2_ solutions for 5 days. Plant phenotype was photographed and recorded, and measurements of plant height, root length, and biomass were taken. To further assess Cd stress on root elongation, 3-day-old seedlings were exposed to a control solution (0.5 mM CaCl_2_) and a treatment solution (0.5 mM CaCl_2_ with 5 µM Cd(NO_3_)_2_) at 30 ℃ for three days. Root lengths were measured at 0, 24, 48, and 72 hours post-exposure. Each rice line and Cd treatment condition had eight biological replicates.

### 2.5. Measurement of Malondialdehyde (MDA)

Lipid peroxidation was assessed by measuring MDA levels with thiobarbituric acid (TBA) assay. Fresh leaf samples (0.6 g) were homogenized in 6 mL of 0.1% (w/v) trichloroacetic acid (TCA). The homogenate was centrifuged at 4500×g for 8 minutes. A 3 mL portion of the supernatant was mixed with 3 mL of 0.6% TBA and incubated in boiling water for 10 minutes. After rapid cooling on ice, the mixture was centrifuged again. The absorbance of the supernatant was measured at 450 nm, 532 nm, and 600 nm, and MDA concentration was calculated according to the TBA reaction as described by Hu et al., 2017.

### 2.5. Soil pot experiment

A soil sample contaminated with Cd was collected from a paddy field located in Xiangtan, Hunan Province, China, exhibiting a total Cd content of 1.13 mg/kg. The collected soil was air-dried, ground, and passed through a 50-mesh sieve to ensure homogeneity.Subsequently, the sieved soil was uniformly blended with fertilizers at a ratio of 0.3 g KH_2_PO_4_ and 0.5 g (NH_4_)_2_SO_4_ per kilogram of soil in a single mixing step. Plastic buckets, each with a 22-liter capacity, were filled with 17 kg of the prepared soil-fertilizer mixture. Tap water was added to each bucket to submerge the soil, which was then allowed to soak for a period of 7 days. The pH of the soil was determined to be approximately 5.7. One seedling from each of the following rice lines - WT plants, the *osnac5* mutant, and two *osnac5* complementation lines - were transplanted into individual plastic buckets. Throughout the vegetative growth phase, the plants were irrigated twice daily to maintain a water level of 1—3 cm above the soil surface. During the rice grain-filling stage, the water was drained from the buckets. Upon reaching grain maturity, the plants were harvested. The biomass and metal concentrations in the grains were determined.

### 2.6. Elemental Concentration Measurement

To evaluate the translocation efficiency of Cd from roots to shoots, 14-day-old plants of WT, the osnac5 mutant line and two osnac5 complemental lines were subjected to 0.1, 1, or 5 µM Cd(NO3)2 for a duration of 7 days. Post-exposure, the roots were meticulously washed twice with 5 mM CaCl2 to remove surface-bound Cd, followed by a thorough rinse with deionized water.The roots and shoots were then separated, harvested and oven-dried at 75 ℃ for 5 days to achieve a constant dry weight. For the determination of metal element content, the dried plant tissues were digested using a mixed acid protocol.Each sample, consisting of a minimum of 0.1 g of dry tissue, was digested in a heating block with a mixture of perchloric acid (HClO4) and nitric acid (HNO3) in a 13:87 (v/v) ratio. The digestion process involved a temperature gradient: the samples were heated at 110 ℃ for 2 hours, followed by 140 ℃ for 1.5 hours, then 160 ℃ for 0.5 hours, and finally maintained at 180 ℃ until complete dryness was achieved.After digestion, the residue was dissolved in 2.5% HNO3 for analysis. The concentration of metal element in the digested samples was quantified using inductively coupled plasma mass spectrometry (ICP-MS, Perkin-Elmer NexION 300x, Waltham, MA, USA).

### 2.7. Chromatin Immunoprecipitation Quantitative PCR assay (ChIP-qPCR)

ChIP-qPCR assays were performed using a commercial chromatin immunoprecipitation Kit (Merck Millipore) in accordance with the method described by Hu et al. (2021). In brief, 5 g of fresh roots harvested from OsNAC5Pro::NAC5-HA transgenic seedlings were subjected to cross-linking with 1.2% formaldehyde under vacuum for 8 minutes.Subsequent to fixation, chromatin was extracted and sonicated to generate fragments ranging from 200 to 1000 base pairs (bp) in length, as per the manufacturer’s instructions.

A total of 100 µL of the sonicated chromatin solution was then immunoprecipitated with 5 µL of anti-HA antibody (Santa Cruz Biotechnology) under gentle agitation at 50 rpm and 4 ℃ overnight. The immunocomplexes were processed to release and purify the DNA fragments associated with the OsNAC5 protein, which were then utilized as templates for qPCR analysis.Genomic fragments from regions not expected to bind OsNAC5 served as negative controls. Quantification of the relative abundance of target DNA sequences was performed using the ΔΔCt (cycle threshold) method.Actin was used as the internal reference gene, and its expression in the ‘no antibody’ (noAB) samples was used as the baseline for normalization.

### 2.8. Electrophoretic mobility shift assay (EMSA)

The *OsNAC5* gene was PCR-amplified from ‘ZH11’ cDNA using Phanta Super-Fidelity DNA Polymerase (Vazyme Biotech Co., Ltd). The resulting PCR product was cloned into the pGEX-4T-1 vector utilizing ClonExpress II One Step Cloning Kit (Vazyme) according to the manufacturer’s instructions. The recombinant GST-OsNAC5 fusion protein was subsequently purified with the MagneGST™ Protein Purification System (Promega), following the provided protocol.

For EMSA, a DNA fragment encompassing the “CATGTG” motif within the OsNramp1 promoter was synthesized and labeled at 5’ end with biotin. Additionally, an unlabeled version of the same DNA fragment was synthesized to function as a competitor in the binding assay (Supplementary Table S1).

### 2.8. Transient Luciferase (LUC) Reporter Assay

The 1214 bp promoter region of *OsNramp1* was amplified using specific primers (Supplemental Table S1). This fragment was cloned into the pCam1381-LUC binary vector to create proNramp1::LUC via a single BamH I restriction site. The construct, along with pCam1300-OsNAC5, was then introduced into A. tumefaciens (strain EHA105). Transient luciferase reporter assays were performed as described by Hu et al., 2021.

### 2.12 Statistical analysis

To analyze the data, we used two-way ANOVA with Tukey’s HSD post hoc tests and Student’s *t* tests when applicable. These analyses were conducted using IBM SPSS Statistics version 25.0. Results are shown as means ± standard deviation, with different letters indicating significant differences (p ≤ 0.05). Significance levels are marked as *p ≤ 0.05, **p ≤ 0.01, and ***p ≤ 0.001.

### 2.9. Gene numbers

*OsNAC5* (LOC_Os11g08210), *Actin* (LOC_Os03g50885), *OsNRAMP1* (LOC_Os07g15460), *OsNRAMP5* (LOC_Os07g15370), *OsHMA2* (LOC_Os06g48720), *OsHMA3* (LOC_Os07g12900), *OsLEA3* (LOC_Os05g46480), *OsYSL2* (LOC_Os02g43370), and *OsNAS3* (LOC_Os07g48980).

## 3. Results

### 3.1. Sequence and subcellular localization of Analysis of OsNAC5

The full-length open reading frame (ORF) of *OsNAC5* was cloned from rice root cDNA, consistent with the sequence in the Rice Annotation Project Database. Structural analysis identified three exons and two introns in *OsNAC5* gene, which encodes a protein of 329 amino acids with a molecular weight of approximately 35.53 kDa. Phylogenetic analysis placed OsNAC5 in the rice NAC transcription factor family, noted for a conserved NAM domain involved in DNA binding (Supplementary Fig. S1A). OsNAC6, the closest homolog, shares 65.23% sequence identity with OsNAC5 (Supplementary Fig. S1B). Previous studies have shown that OsNAC6 responds to stress factors like drought, abscisic acid (ABA), and jasmonic acid (JA), and its overexpression enhances drought and high salinity tolerance in rice (Nakashima et al., 2007; Lee et al., 2017; Rachmat et al., 2014).

Bioinformatic analysis using ProtComp 9.0 program (http://www.softberry.com/) predicted that OsNAC5 is a nuclear protein with a nuclear localization signal (NLS) peptide. This was experimentally confirmed through a transient expression assay in rice protoplasts using an OsNAC5-YFP (yellow fluorescent protein) fusion construct. Confocal laser scanning microscopy showed that the YFP-tagged OsNAC5 colocalized with the red fluorescent protein-tagged Histone 2A (RFP-H2A) nuclear marker (Hu et al., 2017). In contrast, YFP alone was distributed throughout the cell (Fig. 1). These findings align with previous research showing nuclear localization of OsNAC5-GFP in rice roots (Takasaki et al., 2010). Collectively, these results confirm that OsNAC5 is localized in the nucleus.

**Fig. 1.**
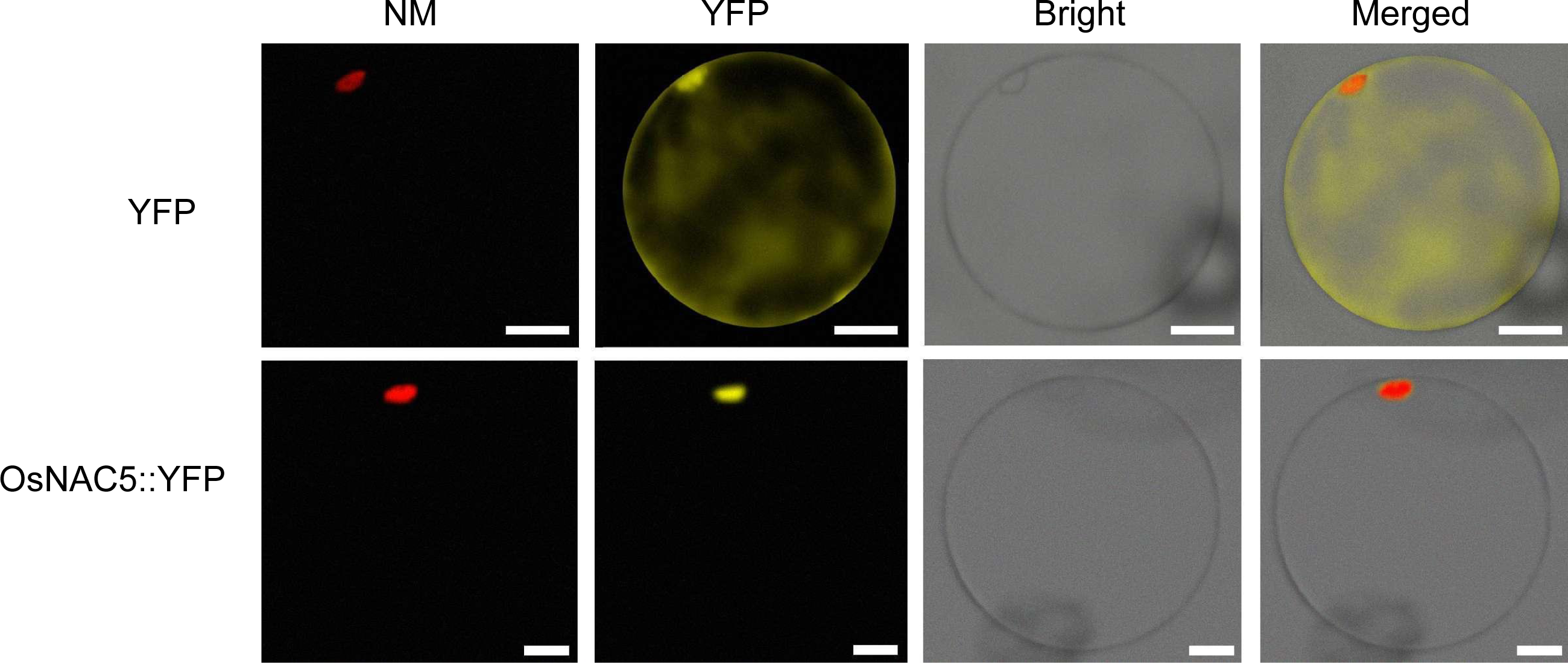
Subcellular localization of OsNAC5. Confocal images of rice protoplast expressiong yellow fluorescent protein gene (YFP) or *OsNAC5-YFP*. NM (nuclear marker) is a nuclear localization tag, red fluorescent protein (RFP) linked to Histone 2A (RFP-H2A). Scale bars = 20 µM.

### 3.2. Expression pattern of OsNAC5

The expression pattern of *OsNAC5* was examined by RT-qPCR in various tissues of hydroponically grown rice plants (cv. ZH11). The analysis showed that *OsNAC5* was mainly expressed in the roots (Fig. 2A). This observation contrasts with Song et al. (2011), who reported that *OsNAC5* was primarily expressed in the leaves of cv. Zhonghua10, indicating that the expression of *OsNAC5* varies among cultivars.

**Fig. 2.**
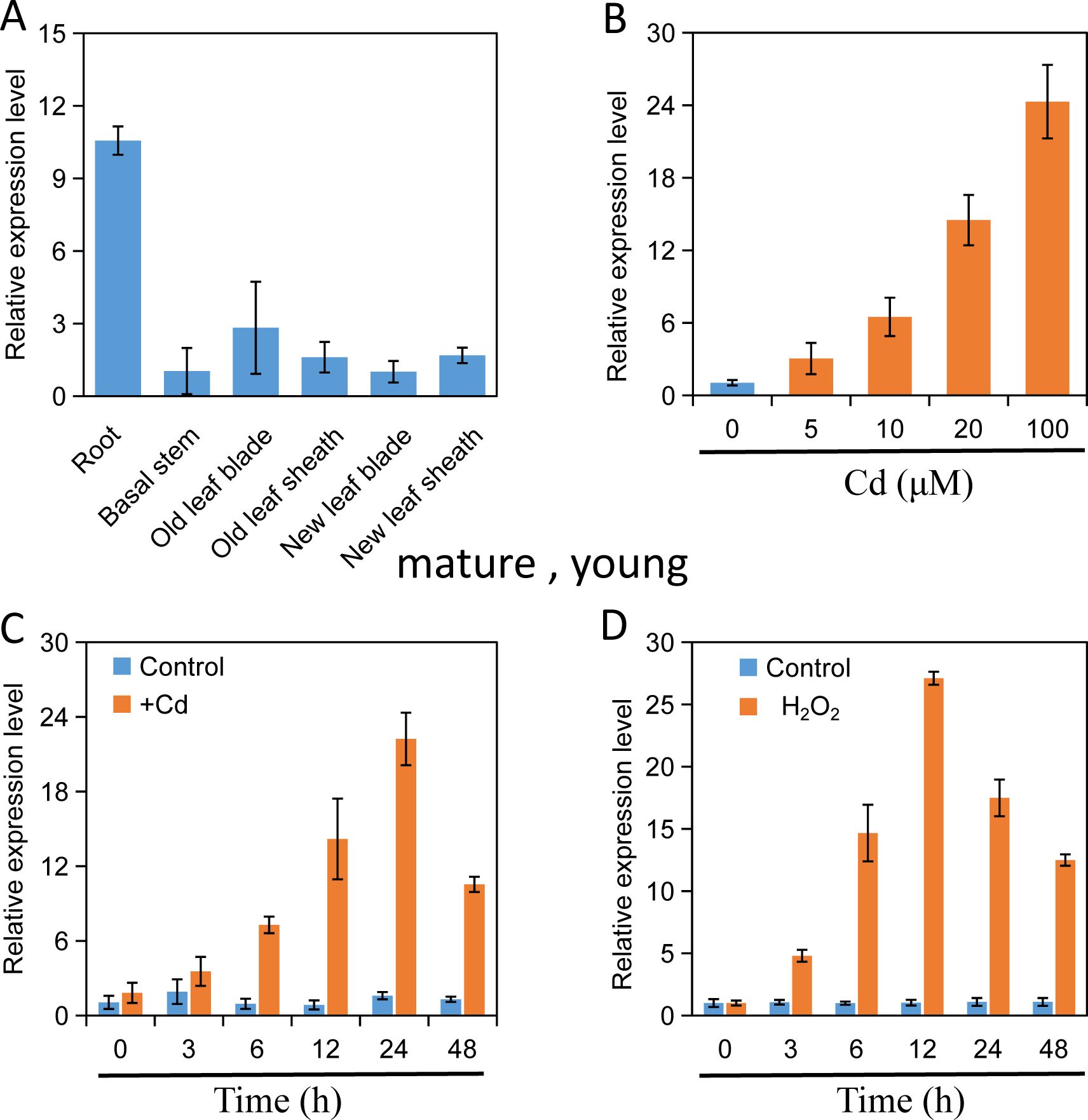
Gene expression pattern of *OsNAC5*. (A) Relative expression of the *OsNAC5 gene* in various tissues of the 2-week-old wild-type rice seedlings. (B) Dose-dependent expression of the *OsNAC5* gene in the roots of wild-type rice in response to Cd stress. Rice seedlings were exposed to a Cd(NO_3_)_2_ solution with varying concentrations (0, 5, 10, 20, 100 μM) of for 24 hours. (C, D) Time-dependent expression of the *OsNAC5* gene in the roots of wild-type rice in response to Cd stress or H_2_O_2_. Rice seedlings were exposed to a solution containing 100 μM d(NO_3_)_2_ 10 mM H_2_O_2_ (to induce stress) for different durations. The expression of the *OsNAC5* gene was determined using RT-qPCR. *Actin* was used as an internal standard. The expression levels were normalized to seedlings without the addition of Cd or H_2_O_2_. Data are represent the means ±standard deviation (SD) of four biological replicates.

To explore the response of *OsNAC5* to Cd or hydrogen peroxide (H_2_O_2_) stress, hydroponic experiments were conducted. Cd exposure led to a dose-dependent increase in *OsNAC5* expression in rice roots (Fig. 2B). A time-course analysis of Cd-treated roots revealed a rapid rise in *OsNAC5* transcript levels, peaking at 24 hours post-treatment (Fig. 2C). Previous research has shown that *OsNAC5* is inducible by various abiotic stresses such as drought, high salinity, and cold (Takasaki et al., 2010; Jeong et al., 2013; Song et al., 2011). Considering the role of H_2_O_2_ as a crucial signaling molecule (Mittler et al., 2011; Ni et al., 2019), we also assessed its effect on *OsNAC5* expression. The results demonstrated a rapid up-regulation of *OsNAC5* following H_2_O_2_ treatment, with peak expression levels at 12 hours (Fig. 2D), suggesting a quicker response to H_2_O_2_ than to Cd stress.

To explore the potential role of H_2_O_2_ in the Cd-induced *OsNAC5* expression pathway, rice plants were pre-treated with two reactive oxygen species (ROS) modulators, dimethylthiourea (DMTU), an H_2_O_2_ scavenger, and diphenyleneiodonium (DPI), an NADPH oxidase inhibitor, before Cd exposure (Zhang et al., 2014). These pre-treatments significantly reduced the Cd-induced increase in *OsNAC5* expression (Supplementary Fig. S2). Indicating that intracellular H_2_O_2_ is a crucial mediator for the Cd-induced up-regulation of *OsNAC5* transcript levels.

### 3.3. Knockout of OsNAC5 results in a Cd-sensitive phenotype in rice

To clarify the physiological role of *OsNAC5* in rice’s response and tolerance to Cd stress, we generated an *osnac5* mutant in the ZH11 background using the CRISPR/Cas9 system. Sanger sequencing confirmed a homozygous *osnac5* mutant line with a 1 bp thymine insertion causing a frameshift mutation at the target gRNA site (Supplementary Fig. S3). To verify the function of *OsNAC5*, we conducted a genetic complementation assay by introducing *OsNAC5* genomic sequence (ProNAC5::NAC5-HA) into the *osnac5* mutant. This resulted in the generation of two complementation lines (C1 and C2), which were confirmed through immunoblotting analysis with anti-HA antibody. These lines were then used for subsequent phenotypic analysis and chromatin immunoprecipitation (ChIP) assay (Supplementary Fig. S4).

Under control conditions, no noticeable differences were observed among WT, the *osnac5* mutant, complementation lines C1 and C2 at two-week-old plants (Figure 3A). However, when exposed to 10 µM Cd(+Cd)or 100 µM Cd(++Cd)for 5 days, the *osnac5* mutant plants showed more severe growth inhibition and leaf necrosis, unlike WT and complementation lines (Figure 3B,C). We measured shoot height, root length, biomass, and MDA concentration under these conditions. Under normal conditions, there were no visible differences in these parameters among the WT, the *osnac5* mutant, and complementation lines (Figure 3D-I). However, under +Cd or ++Cd conditions, the shoot height, root length, and biomass of the *osnac5* mutant were lower than those of the WT and complementation lines. Additionally, the MDA concentration in the shoots and roots of the *osnac5* mutant was higher (Figure 3D-I).

**Fig. 3.**
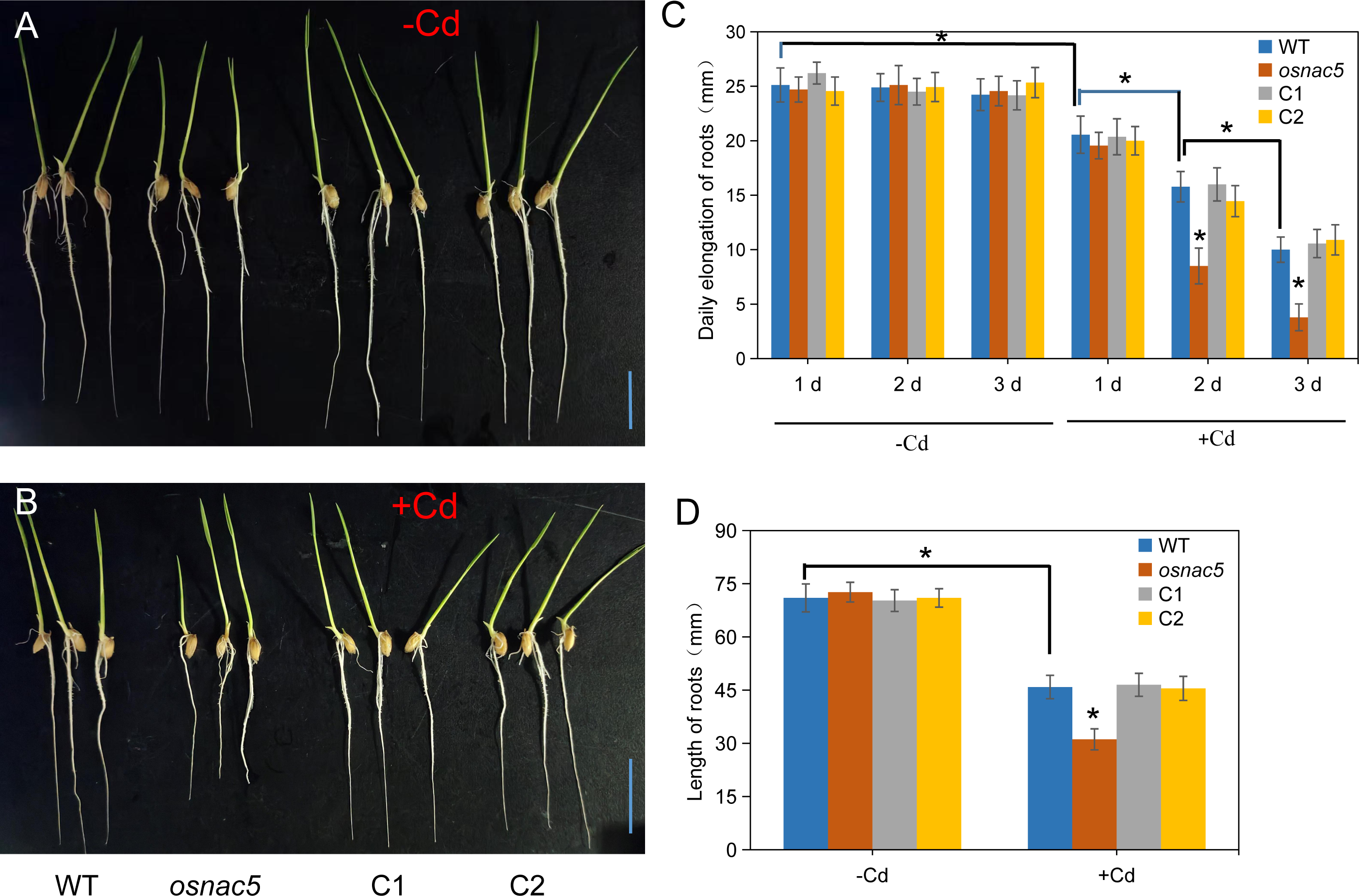
The root elongation of *osnac5* mutant is hypersensitive to Cd stress. (A, B) The growth phenotypes of rice seedling roots treated for 3 days in solutions containing 0.5 mM CaCl₂ (-Cd) or 0.5 mM CaCl₂+5 μM Cd(NO_3_)_2_ (+Cd), respectively. (C) Daily elongation rate of the roots of rice seedlings under - C d and + C d conditions. (D) Length of rice seedling roots after 3 days under -Cd and +Cd conditions. WT refers to wild-type rice, *osnac5* denotes rice with the *OsNAC5* gene knockout line, and C1 and C2 represent complemented lines of *osnac5* mutant line. Mean values ± SD are shown. The values of the indicated genotypes were compared to that of the wild-type (WT) plants (Student’s *t* test, * P < 0.05, n = 8). Bar=20 mm.

Further investigation showed that under normal conditions, the root length of the *osnac5* mutant seedlings was similar to WT (Supplementary Fig. S5A). Under 5 µM Cd for 3 days, the root length of mutants was significantly reduced compared to WT, but complementation lines maintained root length similar to WT under both conditions (Supplementary Fig. S5B). Daily root elongation was about 25 mm for all lines under normal conditions, but Cd stress inhibited elongation on the first day across all lines. On the second and third days, *osnac5* mutants showed more pronounced inhibition compared to WT and complementation lines (Supplementary Fig. S5C). After 3 days of Cd treatment, WT rice had a 36.5% inhibition rate of root elongation compared to control, while the *osnac5* mutants showed a lower rate of 57.1% (Supplementary Fig. S5D). These findings strongly indicate that *OsNAC5* plays a crucial role in the response of rice roots to Cd stress, as its disruption leads to increased Cd sensitivity.

### 3.4 Knockout of OsNAC5 decreased Cd uptake in rice

To explore the role of *OsNAC5* in mediating Cd accumulation in rice, two-week-old plants of WT, the *osnac5* mutant, two complementation lines C1 and C2 were subjected to 0.1, 1 or 5 µM Cd(NO_3_)_2_ for 7 days. At Cd concentration of 0.1 μM, the *osnac5* mutant plants showed a marginal decrease in Cd accumulation in both the roots and shoots compared to the WT and complementation lines, however, this reduction was not statistically significant (P>0.05; Fig. 4A, B). In contrast, at 1 μM Cd exposure, the *osnac5* mutant plants exhibited a significant reduction in Cd concentration by 15.6% in roots and 15.2% in shoots relative to the WT and two complementation lines (both P < 0.05; Fig. 4A, B). This trend persisted at the higher concentration of 5 μM Cd, with the *osnac5* mutants displaying a significant decrease of 15.2% in root Cd concentration and 15.1% in shoot Cd concentration compared to the WT and complementation lines (both P < 0.05; Fig. 4A, B).

**Fig. 4.**
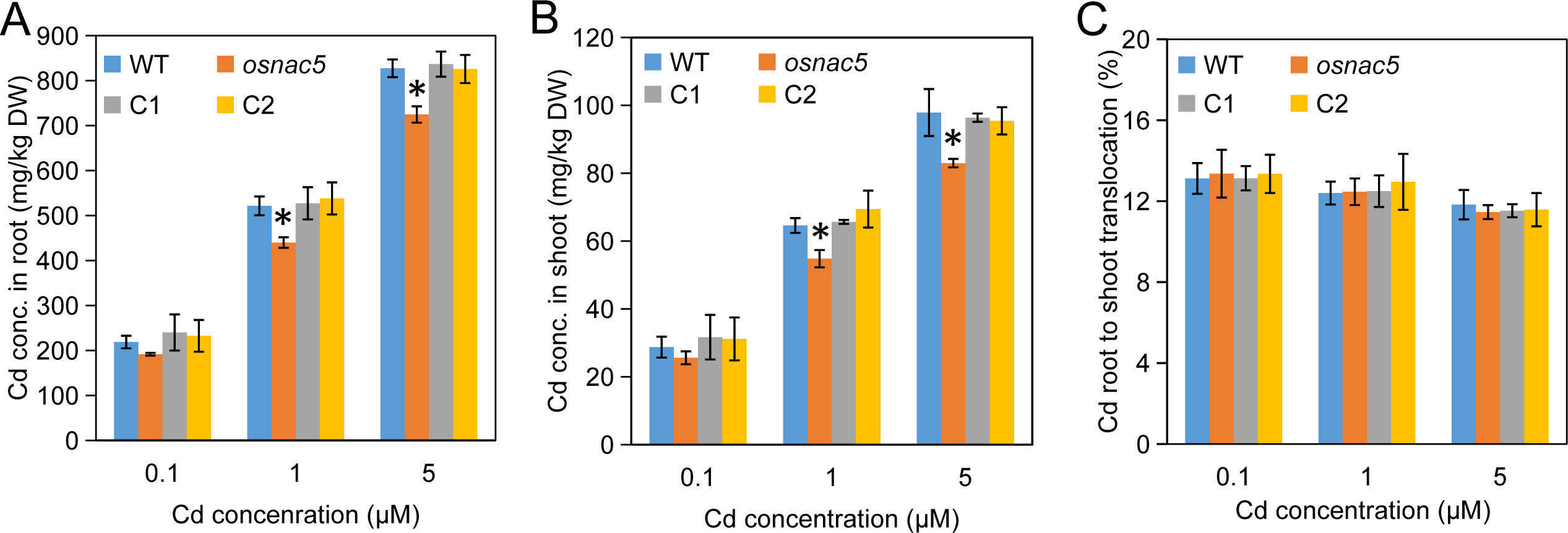
Effect of *OsNAC5* knockout on rice Cd accumulation. The *OsNAC5* knockout line (*osnac5*), two complemental lines (C1 and C2) and wild type (WT, cv. ZH11) were exposed to 0.1, 1 or 5 µM Cd(NO_3_)_2_ for 7days. (A,B) Concentration of Cd in the roots (A) and the shoots (B). (C) The percentage of Cd distributed to the shoot. Roots and shoots were harvested separately and subjected to determination of Cd by ICP-MS. All data were based on comparisons with WT plants. DW: dry weight. Values represent means ± SD of biological replicates (n = 4). Statistical comparison was performed by Tukey’s test (*P<0.05).

Conversely, the Cd concentrations in the shoot and root tissues of the complementation lines did not differ significantly from those of the WT under any of the tested Cd treatments (Fig. 4A, B). Additionally, the efficiency of Cd translocation from roots to shoots did not show significant variation among the WT, the *osnac5* mutant, and complementation lines under three Cd treatment conditions (Fig. 4C). These results indicate that the *osnac5* knockout may lead to reduced Cd uptake in rice without significantly altering the translocation of Cd from roots to shoots.

### 3.5 OsNRAMP1 is induced by Cd stress dependent on OsNAC5

To further examine the potential link between *OsNAC5* knockout and Cd accumulation in rice, we performed real-time qPCR analysis of these Cd transporter genes. After 12 hours of exposure to 10 μM Cd, the expression of *OsNRAMP1* in WT rice roots was significantly upregulated, showing an approximate 5.6-fold increase compared to the untreated control (Fig. 5A). In contrast, the *osnac5* mutant did not exhibit any change in *OsNRAMP1* expression levels upon Cd treatment (Fig. 5A). The complementation lines C1 and C2, however, showed a strong induction of *OsNRAMP1* expression, mirroring the response seen in the WT rice (Fig. 5A).

**Fig. 5.**
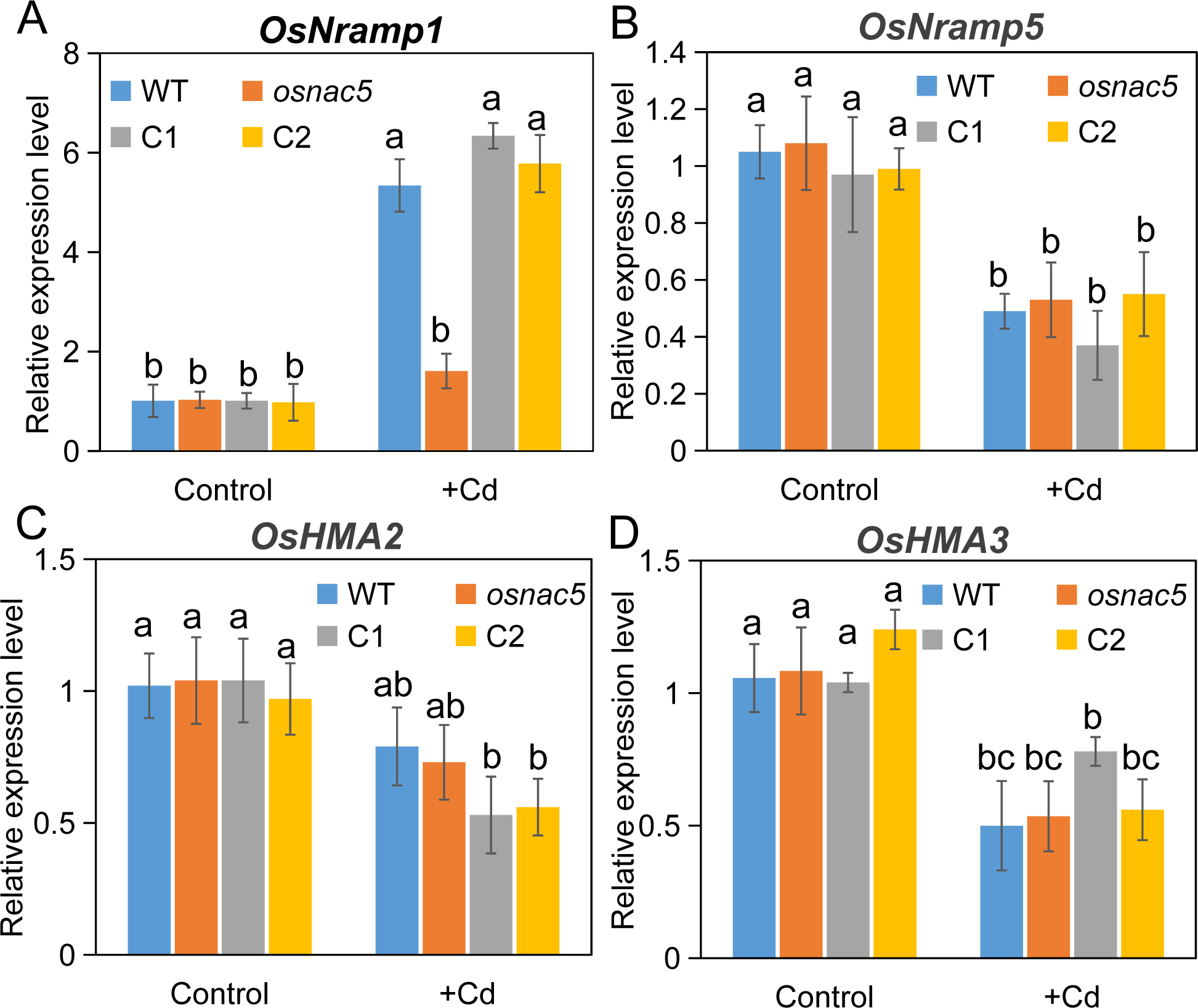
Transcript levels of Cd transporter genes *OsNramp1* (A), *OsNramp5* (B), *OsHMA2* (C) and *OsHMA3* (D) in roots of Cd-treated rice plants. Two-week-old wild-type rice seedlings were exposed to 0.1 µM Cd(Control) or 10 µM Cd(+Cd) for 12 h. Real-time qPCR analysis was used to determine the relative transcript levels shown. *Actin* was used as internal standards. Values are mean ± SD (n = 4). Different letters above bars indicate significant differences among rice plants lines under control or +Cd treatment (P<0.05 by Tukey’s test).

Under the same Cd exposure, the expression levels of *OsNRAMP5* and *OsHMA3* in WT rice roots were significantly downregulated to about 0.5-fold of the control, whereas *OsHMA2* expression remained unchanged (Fig. 5B–D). Notably, there were no significant differences in the expression levels of *OsNRAMP5*, *OsHMA2*, and *OsHMA3* between the WT, the *osnac5* mutant, and complementation lines (Fig. 5B–D). Additionally, under control conditions, the transcript levels of these four Cd transport-related genes were similar across the roots of WT, the *osnac5* mutant, and complementation lines (Fig. 5A–D). These observations suggest that of the four Cd transporters analyzed, only *OsNRAMP1* expression is induced by Cd stress in a manner dependent on the functional presence of *OsNAC5*.

We also assessed whether the *OsNAC5* knockout would alter the gene’s own expression response to Cd stress. Real-time qPCR analysis indicated that under Cd stress, the expression level of *OsNAC5* in the *osnac5* mutant was similar to that in the WT rice (Supplementary Fig. S6A). Given that the reduced Cd accumulation in the *osnac5* mutant does not account for its increased sensitivity to Cd stress, it is postulated that *OsNAC5* may regulate the expression of other stress-related genes. The “late embryogenesis abundant” gene *OsLEA3*, known to be directly upregulated by OsNAC5 and involved in various stress responses (Takasaki et al., 2010), was found to be strongly induced by Cd stress in an OsNAC5-dependent manner (Supplementary Fig. S6B). This may partly explain the *osnac5* mutant’s heightened sensitivity to Cd stress. Furthermore, Jeong et al. (2013) showed that overexpression of *OsNAC5* enhances the expression of genes involved in the homeostasis of divalent metals, such as *OsNAS3* and *OsYSL2*. In our study, we found that *OsNAS3* expression was suppressed by Cd stress, but it was unaffected by the *OsNAC5* knockout (Supplementary Fig. S6C). On the other hand, *OsYSL2* expression was not influenced by either Cd stress or OsNAC5 status (Supplementary Fig. S6C). These findings suggest that OsNAC5’s biological function is multifaceted, encompassing the regulation of Cd transport as well as broader stress response mechanisms.

### 3.6 OsNAC5 binds directly to the promoter of OsNramp1

The binding of transcription factors (TFs) to cis-regulatory elements in gene promoters is crucial for gene expression regulation. OsNAC300 has a strong affinity for the NAC transcription factor binding element motif (CATGTG) (Hu et al., 2021).In the ∼1.0 kb *OsNRAMP1* promoter region, two “CATGTG” motifs were identified at −846 to −841 bp and −131 to −126 bp upstream of the transcription start site (Fig. 6A). To examine if OsNAC5 interacts with these motifs in vivo, ChIP-qPCR assays were performed on ProOsNAC5::OsNAC5-HA transgenic rice using an anti-HA monoclonal antibody.Specific promoter region amplicons (S1, S2, and S3) were enriched. Fragments S1 and S3 containing “CATGTG” motifs showed significant enrichment compared to controls, with Cd exposure resulting in higher enrichment than in the absence of it (Fig. 6B). To verify OsNAC5’s binding affinity to the “CATGTG” motif in vitro, EMSA was conducted using recombinant GST-OsNAC5 fusion protein and synthetic oligonucleotides from *OsNRAMP1* promoter. The EMSA results demonstrated a strong DNA binding complex with the “CATGTG” motif, confirming OsNAC5’s specific interaction with the motif in vitro (Fig. 6C). These findings indicate that OsNAC5 directly binds to “CATGTG” motifs in *OsNRAMP1* promoter, regulating gene expression in response to Cd stress.

**Fig. 6.**
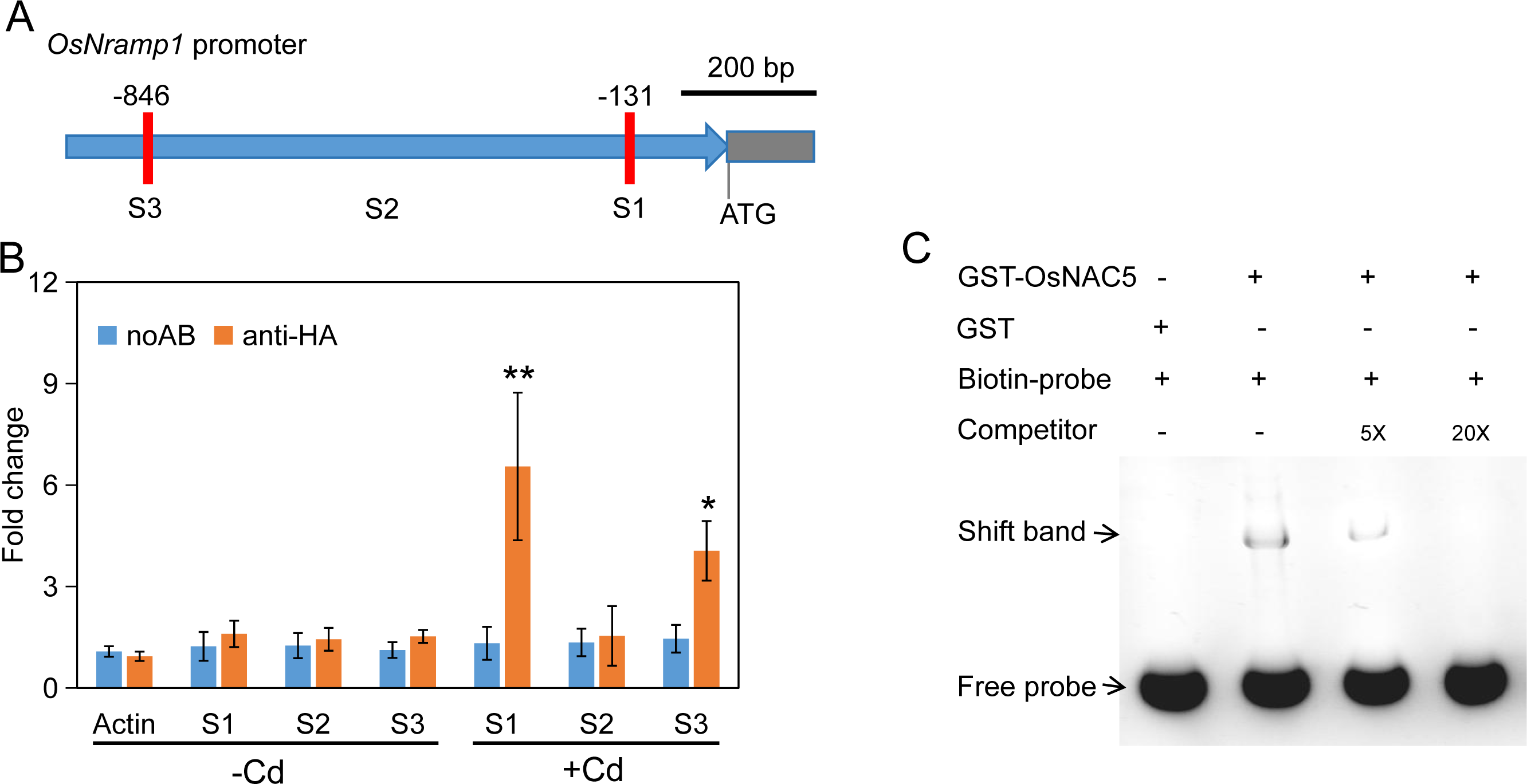
OsNAC5 binds directly to the promoter of *OsNramp1.* (A) Diagram of *OsNramp1* promoter with two “CATGTG” motifs. Red vertical bars represents “CATGTG” motif. Site 1 and site2 indicates genomic DNA fragments around *OsNramp1* promoter for ChIP-qPCR. (B) ChIP-qPCR assay. Binding of OsNAC5 protein to specific regions of the *OsNramp1* promoter was examined with 2-week-old *OsNAC5Pro::NAC5-HA* transgenic seedlings grown to 1/2 Kimura (-Cd) and 1/2 Kimura 100 μM Cd (+Cd) for 6 h. The noAB (no antibody) was used as the control and *Actin* in ‘noAB’ samples was used as the internal standard. Data are means ± SD (*n* = 4). Statistical comparison between noAB and anti-HA was performed by Student’s *t* test. Asterisks indicates significant differences from the control (\**p* ≤ 0.05, \*\**p* ≤ 0.01). ChIP, chromatin immunoprecipitation; HA, the human influenza hemagglutinin tag protein; GST, gulathione S-transferase. (C) Electrophoretic mobility shift assay. Biotin-labeled probe contains the “CATGTG” motif in the *OsNramp1* promoter. Competition is unlabeled probe. Black arrows indicate the shifted bands and free probes.

To determine if OsNAC5 activates *OsNramp1* promoter, a luciferase (LUC) assay was conducted in tobacco leaves using the firefly LUC gene as a reporter. *OsNAC5* was placed under the control of the CaMV 35S promoter. Coexpression of 35S::OsNAC5 with proNramp1::LUC increased the fluorescent signal (Fig. 6D) and showed higher LUC relative expression levels than the control (Fig. 6E).This demonstrates that OsNAC5 directly enhances the transcriptional activity of the *OsNramp1* promoter.

### 3.7 Knockout of OsNAC5 reduced Cd accumulation in rice grain

To assess the impact of *OsNAC5* knockout on Cd accumulation in rice grains (unpolished brown rice), a pot experiment was conducted with the WT, the *osnac5* mutant, and complementation lines (C1 and C2) grown in Cd-contaminated paddy soil (total Cd concentration of 1.13 mg/kg, pH 5.77). Upon reaching maturity, no significant differences were observed among the four lines in terms of plant height, effective panicle number per plant, seed-setting rate, or grain yield per plant (Supplementary Table S2). However, the *osnac5* mutant displayed a marked reduction in Cd concentrations, showing a 39.4% decrease in grains and a 41.8% decrease in husks compared to the WT (Fig. 7). In contrast, the Cd concentrations in the grains and husks of the two complementation lines were similar to those observed in the WT (Fig. 7).

**Fig. 7.**
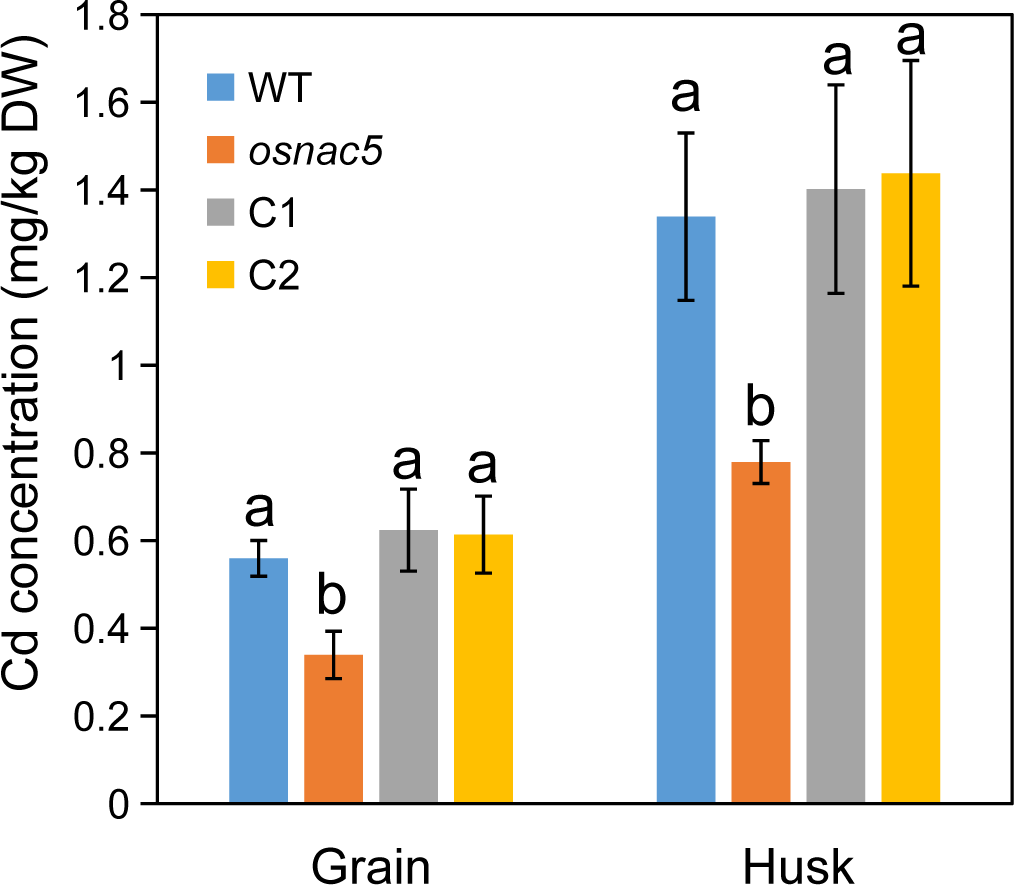
The concentrations of Cd in rice grain (unpolished brown rice) and husk. Wild type (cv. ZH11), *osnac5* mutant and two complementation lines(C1 and C2) grown in a Cd-contaminated paddy soil. Values are mean ± SD (n = 4). Different letters above bars indicate significant differences among rice plants lines (Student’s *t* test, P < 0.05).

Given that OsNRAMP1 is also known to transport Mn (Chang et al., 2020), we measured Mn concentrations in rice grains and detected no significant differences among the four rice lines (Supplementary Fig. S7A). Recent studies have indicated that overexpression of the *OsNAC5* gene enhances the translocation of Fe and Zn from vegetative tissues to seeds (Wairich et al., 2023). Consequently, we measured the concentrations of Fe and Zn in rice grains and found no significant differences among the four lines (Supplementary Fig. S7B, C). These results are consistent with the notion that Fe, Zn, and Mn are essential micronutrients, and plants have developed intricate regulatory mechanisms to maintain their homeostasis, avoiding both deficiency and toxicity.

## 4. Discussion

Previous research has established that *OsNAC5* expression is induced by a variety of adverse environmental stressors, including drought, cold, high salinity, abscisic acid (ABA), and methyl jasmonate (Takasaki et al. 2010; Song et al., 2011). In line with these observations, our study demonstrates that *OsNAC5* is also responsive to heavy metal Cd and H_2_O_2_. Intriguingly, the response to H_2_O_2_ was observed as early as three hours post-treatment, whereas the response to Cd stress was significantly delayed in comparison to H_2_O_2_ (Fig. 2B-D). Moreover, we have shown that the induction of *OsNAC5* by Cd stress is contingent upon intracellular H_2_O_2_ production (Supplementary Fig. S2). These results imply that *OsNAC5* may be implicated in a broad spectrum of biotic and abiotic stress responses, although its specific role in biotic stress remains to be elucidated.

Previous work has shown that overexpression of *OsNAC5* under the maize ubiquitin promoter significantly enhances salt tolerance in rice (Takasaki et al., 2010). Conversely, rice lines with reduced *OsNAC5* expression through RNA interference (RNAi) display decreased tolerance to cold, drought, and high salinity, while overexpression of *OsNAC5* in rice or *Arabidopsis* confers increased resistance to these stressors (Song et al., 2010). Additionally, driving *OsNAC5* expression in rice roots with the root-specific promoter RCc3 leads to increased root diameter, improved drought tolerance, and higher yield (Jeong et al., 2013). In our study, we utilized CRISPR-Cas9 technology to generate an *osnac5* mutant and found that this mutant exhibited increased sensitivity to Cd stress compared to WT plants (Fig. 3B, D). However, no phenotypic differences were detected between the *osnac5* mutant and WT plants under normal growth conditions (Fig. 3A, D).

Both *OsNAC5* and *OsNRAMP1* are induced by Cd stress (Fig. 2B, 5A). Notably, in the *osnac5* mutant, *OsNRAMP1* does not respond to Cd stress induction (Fig. 5A). However, in the complementation lines, *OsNRAMP1* expression is restored to WT levels (Fig. 5A). Interestingly, the loss of *OsNAC5* function does not influence the extent of Cd stress-induced suppression of *OsNRAMP5* expression (Fig. 5B). Considering the established transcriptional activation function of the OsNAC5 protein (Takasaki et al., 2010), our findings indicate that OsNAC5 exerts a distinct positive transcriptional regulatory influence on *OsNRAMP1* in response to Cd stress, while it does not appear to regulate *OsNRAMP5* in the same manner. This specificity underscores the nuanced role of OsNAC5 in the plant’s adaptive response to heavy metal stress, selectively modulating the expression of key transporters involved in Cd uptake.

Moreover, our observations revealed that under non-stress conditions (0.1 µM Cd, serving as the control), the loss of OsNAC5 function does not affect the Cd concentration in the roots and shoots of rice plants (Fig. 4A, B). Correspondingly, the expression level of *OsNRAMP1* is not impacted by the loss of *OsNAC5* function under these conditions (Fig. 5A). These findings provide additional support for the hypothesis that OsNAC5 specifically increases *OsNRAMP1* expression under Cd stress conditions. Chromatin immunoprecipitation followed by quantitative PCR (ChIP-qPCR) experiments confirmed that OsNAC5 interacts more strongly with the NAC binding elements in the *OsNRAMP1* promoter in the presence of Cd stress in vivo (Fig. 6B). Consequently, we propose that OsNAC5 directly associates with *OsNRAMP1* promoter to positively influence its expression during Cd stress, highlighting a targeted regulatory mechanism activated by heavy metal exposure (Fig. 8).

**Fig. 8.**
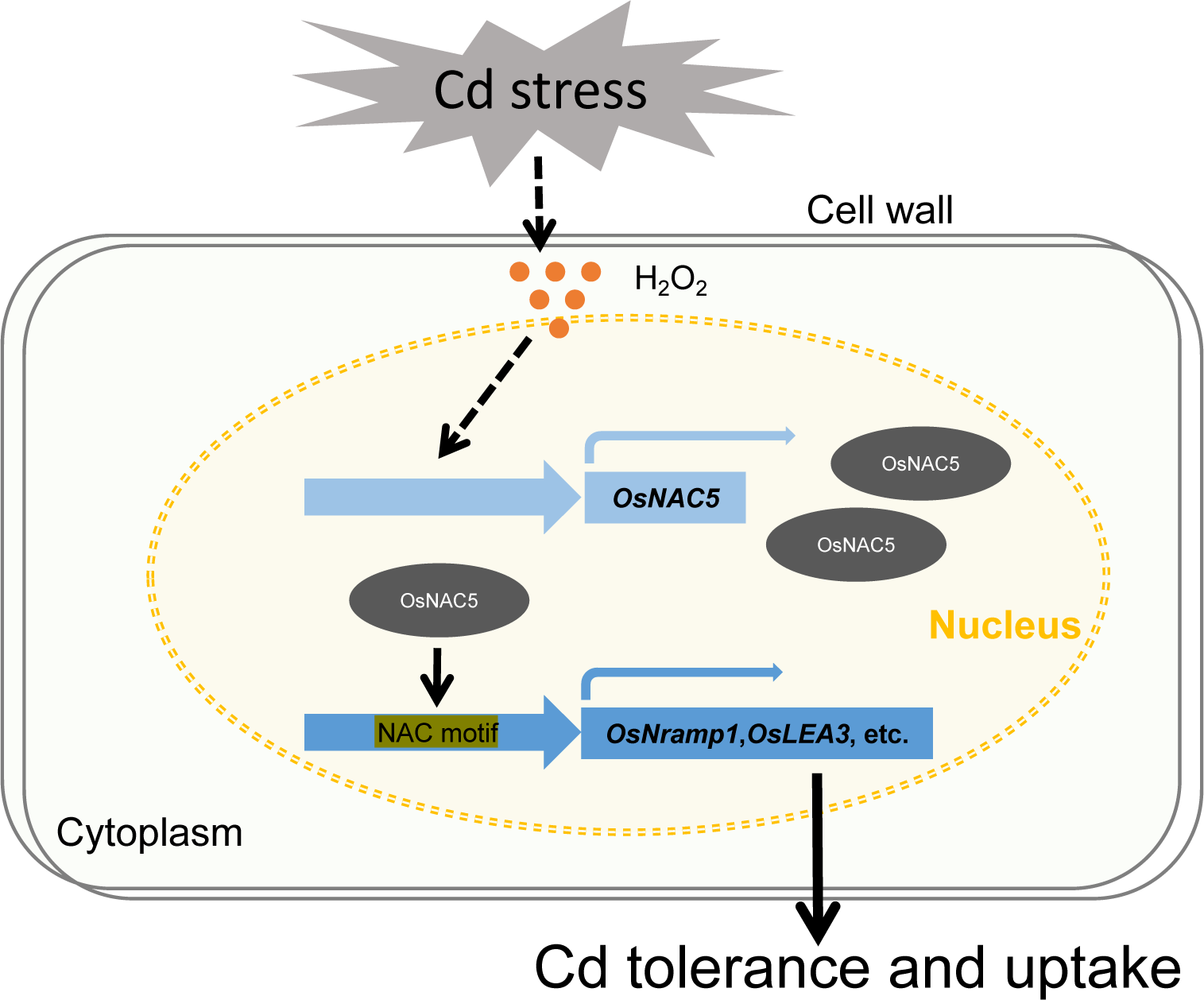
Proposed working model for OsNAC5 regulating of Cd stress response. Cd stress induces an increase in OsNAC5 expression levels, which is dependent on hydrogen peroxide (H_2_O_2_) production triggered by Cd exposure. Subsequently, OsNAC5 directly upregulates the transcription of *OsNramp5* and *OsLEA3*, key genes involved in the cellular response to Cd stress.

OsHMA2, a plasma membrane transporter for Cd, is localized in the pericycle of roots and is constitutively expressed during the vegetative stage (Yamaji et al., 2013). Loss-of-function mutations in OsHMA2 lead to diminished translocation of Cd from roots to shoots, as previously reported (Satoh-Nagasawa et al., 2012; Takahashi et al., 2012; Yamaji et al., 2013). Our study indicates that the expression of *OsHMA2* is minimally influenced by Cd stress and does not appear to be associated with the function of *OsNAC5* (Fig. 4C). OsHMA3, located in the tonoplast of all root cells, has been shown to have its expression modestly reduced by Cd in a cultivar-dependent manner (Ueno et al., 2010; Yan et al., 2016). In line with these findings, we observed that Cd stress suppresses the expression of *OsHMA3*, but this suppression is independent of *OsNAC5* function (Fig. 4D).

Recent molecular biology strategies have aimed to mitigate Cd accumulation in rice grains. CRISPR/Cas9-mediated knockout of *OsNRAMP5* in both japonica and indica rice varieties has been demonstrated to significantly lower Cd accumulation in seeds (Tang et al., 2017; Yang et al., 2019). Overexpression of *OsNRAMP5* has been shown to disrupt radial Cd transport into the stele, thereby reducing Cd accumulation in rice grains (Chang et al., 2020). Similarly, CRISPR/Cas9-mediated knockout of *OsNRAMP1* in japonica rice (ZH11) has been effective in significantly reducing Cd content in both grains and husks (Chang et al., 2020). Alterations in *OsHMA2* expression have also been reported to significantly decrease Cd levels in rice seeds (Takahashi et al., 2012). Moreover, overexpression of *OsHMA3a* from japonica rice can reduce grain Cd content by over 90% without compromising essential nutrient levels such as Zn and Fe (Ueno et al., 2010). In our research, we found that knockout of the transcription factor *OsNAC5* similarly results in reduced Cd content in japonica rice grains (ZH11) (Fig. 7). Unlike the transporters previously mentioned, transcription factors such as OsNAC5 do not directly mediate Cd uptake or transport. This discovery provides an alternative molecular biology strategy to regulate Cd content in rice grains, expanding the toolkit for food safety.

## 4. Conclusion

In addition to its established roles in mediating responses to drought, cold, high salinity, and ABA, our research has unveiled a novel function of the rice transcription factor OsNAC5 in responding to Cd stress. We have demonstrated that OsNAC5 is pivotal in modulating rice’s response to Cd stress and its subsequent accumulation in plant tissues. The expression of the *OsNAC5* gene is upregulated in the presence of Cd stress, a response that is contingent upon the production of H_2_O_2_. Notably, OsNAC5 specifically interacts with the “CATGTG” motifs within the promoter region of *OsNRAMP1*, thereby orchestrating its expression in response to Cd exposure (Fig. 8). Crucially, the knockout of *OsNAC5* leads to a reduction in Cd accumulation in rice grains, underscoring its potential utility in mitigating heavy metal content in crops. These insights not only enhance our comprehension of the molecular mechanisms governing the response to Cd stress in rice but also offer promising avenues for the development of biotechnological strategies to minimize Cd uptake and accumulation in agricultural produce.

## Acknowledgements

This work was supported by the Anhui Provincial University Innovation Team Project, Digital Agriculture Innovation Team (2023AH010039); the National Natural Science Foundation of China (41907145); key project support from the Anhui Provincial Department of Education (2022AH051048, 2023AH050484); a research project of Anqing Normal University (100001195); and the Open Project Program of the Provincial Key Laboratory of the Biodiversity Study and Ecology Conservation in Southwest Anhui Province(Wsz202205,Wxn202303,Wxn202309).

## Environmental Implication

Cadmium (Cd) is readily taken up by plants and can accumulate to toxic levels in edible tissues, posing a significant risk to human health upon consumption. The transporter OsNRAMP1 has been identified as a critical gateway for Cd uptake in plants, yet the intricacies of its regulation have remained largely unexplored. Our research uncovers the pivotal role of the transcription factor OsNAC5 in directly upregulating *OsNRAMP1* expression, thereby modulating Cd uptake and enhancing the plant’s tolerance to Cd stress.

## Supplementary Data

Supplementary Fig. S1. Amino acid sequence multiple alignment and phylogenetic tree analysis were conducted to compare NAC family proteins in rice, Arabidopsis, maize, and wheat.

Supplementary Fig. S2. Cd stress induces the expression of *OsNAC5* in rice roots depending on H_2_O_2_ production.

Supplementary Fig. S3. Identification of the *osnac5* mutant generated by CRISPR/Cas9.

Supplementary Fig. S4. Two complementary lines (ProOsNAC5::OsNAC5-HA) were identified through immunoblotting analysis using an anti-HA antibody.

Supplementary Fig. S5. The root elongation of the *osnac5* mutant is hypersensitive to Cd stress.

Supplementary Fig. S6. Transcript levels of Cd transporter genes *OsNAC5* (A), *OsLEA3* (B), *OsNAS3* (C), and *OsYSL2* (D) were analyzed in the roots of Cd-treated rice plants.

Supplementary Fig. S7. The concentrations of Mn (A), Zn (B), and Fe (C) in unpolished brown rice grains of wild type (cv. ZH11), the *osnac5* mutant, and two complementation lines (C1 and C2) were measured.

Supplementary Table S1.Primers sequence used in this study.

Supplementary Table S2. The analysis of agronomic traits in WT plants (ZH11), the *osnac5* mutant, and two *osnac5* complementation lines (C1 and C2).

