## Supplementary material for "The Transcription Factor OsNAC5 Mediates Cadmium Stress Response and Accumulation in Rice": Supplemental Figures.pdf

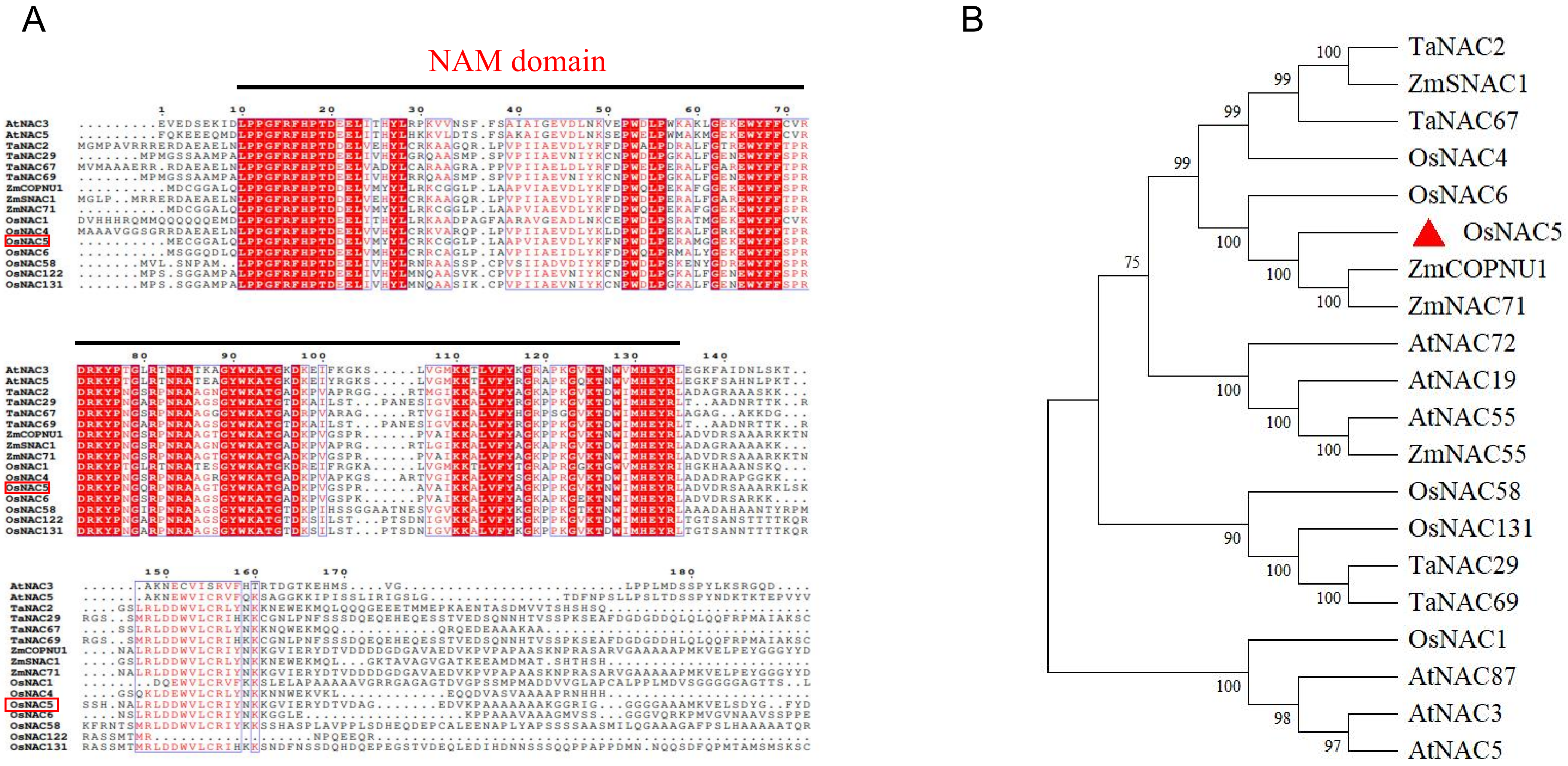


**Supplementary Fig. S1.** Amino acid sequence multiple alignment and phylogenetic tree analysis of NAC family proteins in rice, Arabidopsis, maize, and wheat.
(A) Alignment of sequences using ClustalX and visualization with ESPript 3.x. In the aligned amino acid sequences, red indicates conserved sequences, and blue boxes represent conserved sequences with a global similarity score > 0.7 (threshold input for similarity calculation). The NAM domain is located below the black bar. (B) Phylogenetic analysis of NAC family DNA sequences in rice (Oryza sativa V7_JGI), Arabidopsis (Arabidopsis thaliana TAIR10), maize (Zea mays PH207 v1.1), and wheat (Triticum aestivum Taigu Genotype). Clustering and evolutionary analysis were conducted using MEGA 11 software. The Maximum Likelihood (ML) method was used to construct the phylogenetic tree, the Poisson model was employed to calculate genetic distances, and the Bootstrp method was used to assess tree quality with 1,000 replications. The rice NAC family proteins were used to build this phylogenetic tree using the Maximum Likelihood method with 1,000 bootstrap replicates. The numbers on each branch indicate bootstrap percentages. The red solid triangulum represents OsNAC5.

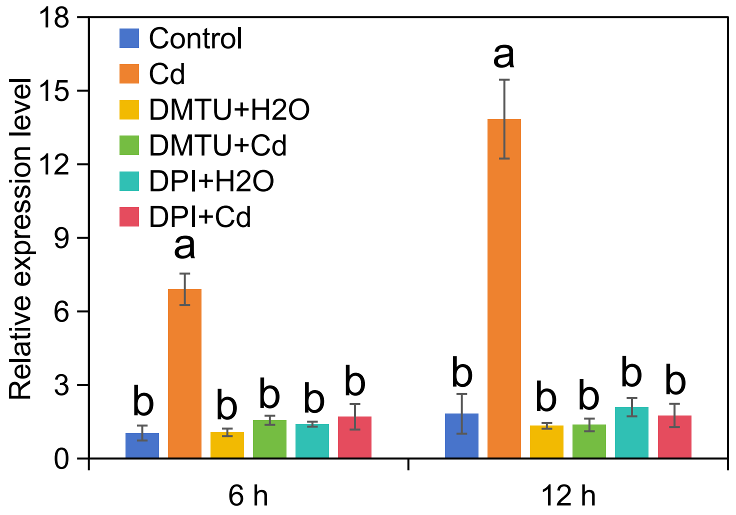


**Supplementary Fig. S2.** Cd stress induces the expression of *OsNAC5* in rice roots depending on $H_2O_2$ production. Effects of pre-treatments with dimethylthiourea (DMTU) and diphenylene iodonium (DPI) on the expression of *OsNAC5* in rice roots exposed to Cd treatment. Two-week-old wild-type rice seedlings were pre-treated with 5 mM DMTU or 100 μM DPI for 2 h, and then exposed to 100 μM Cd for 6 h and 12 h, respectively. Relative expression levels of the *OsNAC5* gene were analysed by RT–qPCR. Values represent means ±SE of three biological replicates. Means indicated by the same letter did not show significant differences ($P<0.05$ by Tukey's test).

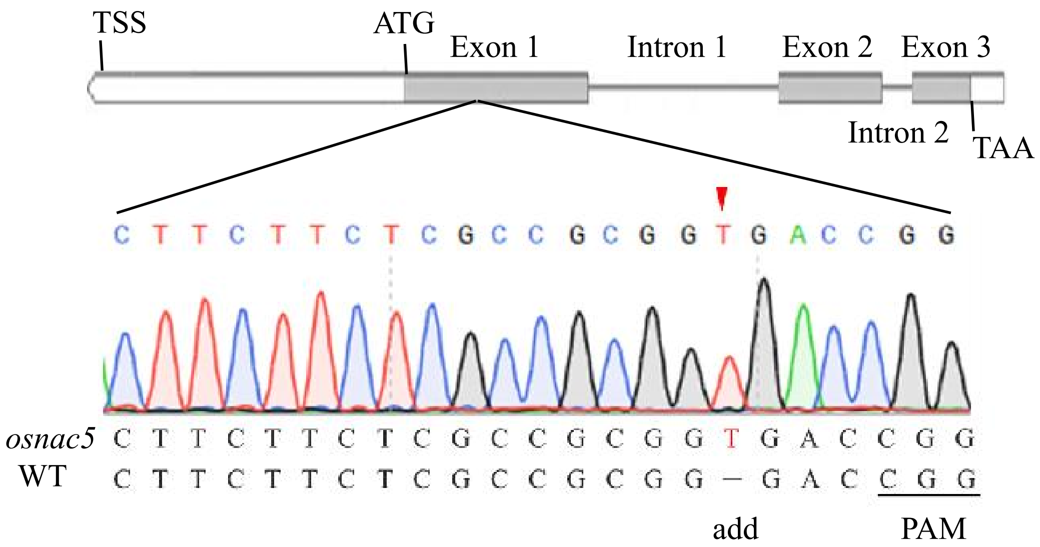


**Supplementary Fig. S3.** Identification of the *osnac5* mutant generated by CRISPR/Cas9.

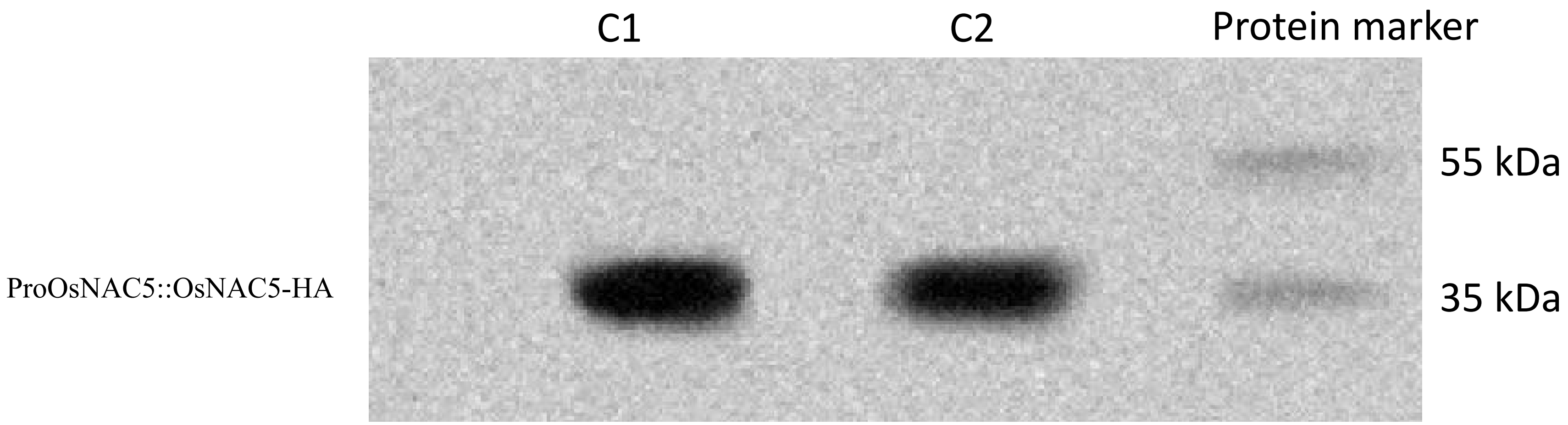


**Supplementary Fig. S4.** Two complementary lines (ProOsNAC5::OsNAC5-HA) were determined by immunoblotting analysis with anti-HA antibody.

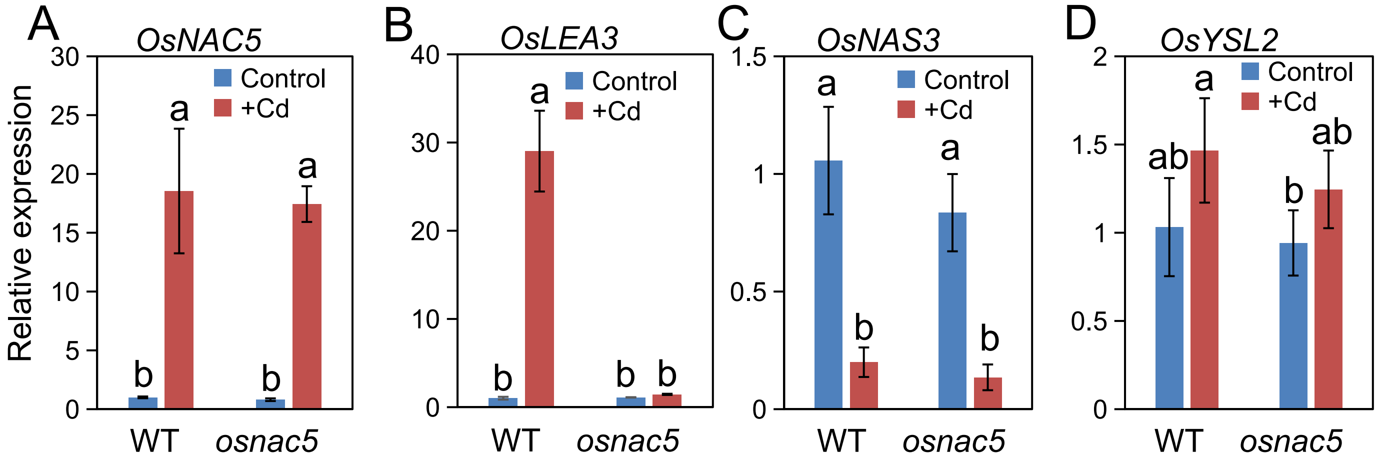


**Supplementary Fig. S5.** Transcript levels of Cd transporter genes *OsNAC5* (A), *OsLEA* (B), *OsNAS3* (C) and *OsYSL2* (D) in roots of Cd-treated rice plants. Two-week-old wild-type rice seedlings were exposed to 0.1 μM Cd(Control) or 10 μM Cd(+Cd) for 12 h. Real time qPCR analysis was used to determine the relative transcript levels shown. *Actin* was used as internal standards. Values are mean ± SD (n = 4). Different letters above bars indicate significant differences among rice plants lines under control or +Cd treatment ($P<0.05$ by Tukey's test).

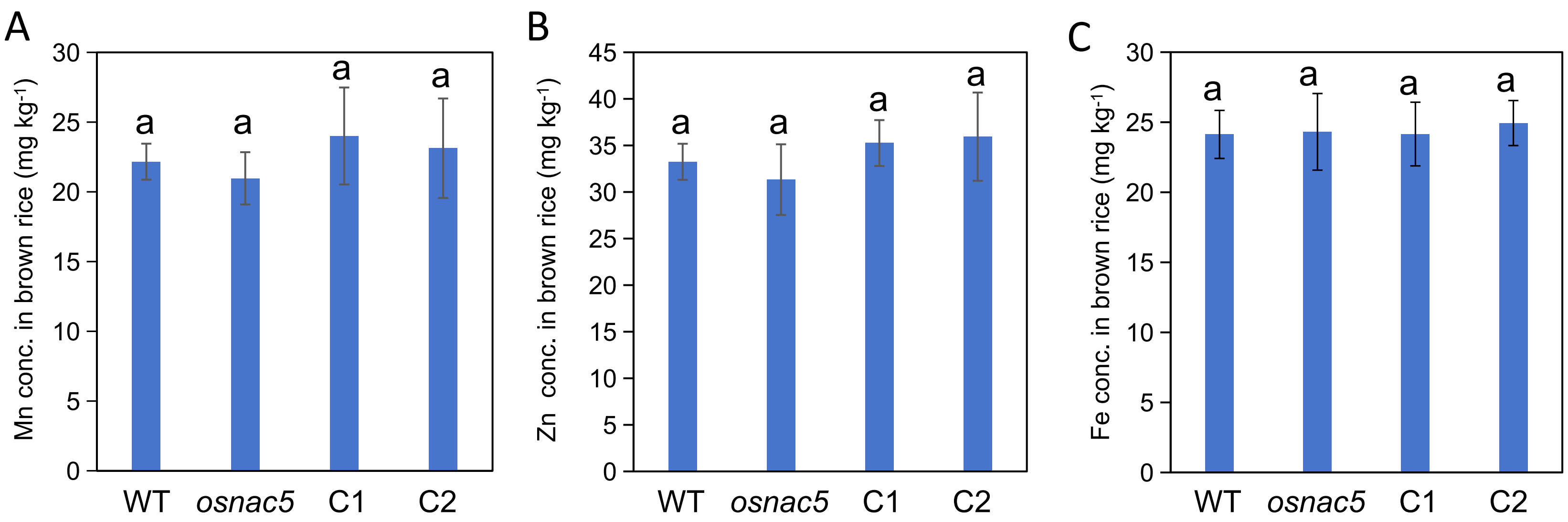


**Supplementary Fig. S6.** The concentrations of Mn (A), Zn (B) and Fe (C) in rice grain (unpolished brown rice) of wild type (cv. ZH11) , *osnac5* mutant and two complementation lines (C1 and C2). Values are mean ± SD (n = 4). Different letters above bars indicate significant differences among rice plants lines at $P<0.05$ ( Student's *t* test).
