## Supplementary material for "The Transcription Factor OsNAC5 Mediates Cadmium Stress Response and Accumulation in Rice": Supplemental Table S2.pdf

Supplementary Tab. S2. Analysis of agronomic traits in WT plants (ZH11), the *osnac5* mutants and two *osnac5* complementation lines (C1 and C2).

| Line | Plant height (cm) | Effective panicle number per plant | Length of main panicle (cm) | Grain number per panicle | Seed-setting rate (%) | Grain yield per plant (g) |
|---|---|---|---|---|---|---|
| WT | 107.2±2.86 a | 21.7±2.37 a | 21.33±1.23 a | 117.00±9.15 a | 82.25±5.59 a | 56.87±9.48 a |
| *osnac5* | 105.4±2.80 a | 21.3±2.90 a | 19.4±1.36 b | 105.20±7.12 a | 79.50±4.15 a | 47.91±8.00 a |
| C1 | 108.5±3.24 a | 23.6±4.21 a | 20.56±2.64 a | 111.25±8.52 a | 80.98±6.24 a | 52.68±10.21 a |
| C2 | 106.3±3.55 a | 22.0±3.45 a | 21.55±1.98 a | 114.62±7.86 a | 78.52±4.59 a | 54.25±9.75 a |

The data in the table are presented as mean values ± standard deviation. Within each column, values accompanied by the same letter are not significantly different, whereas those with different letters are significantly different, as determined by statistical analysis at the 0.05 probability level.
